# Colonic epithelial adaptation to EGFR-independent growth induces chromosomal instability and is accelerated by prior injury

**DOI:** 10.1101/2021.02.02.429426

**Authors:** Tiane Chen, Maged Zeineldin, Blake Johnson, Yi Dong, Akshay Narkar, Taibo Li, Jin Zhu, Rong Li, Tatianna C. Larman

## Abstract

Although much is known about the gene mutations required to drive colorectal cancer (CRC) initiation, the tissue-specific selective microenvironments in which neoplasia arises remains less characterized. Here, we determined whether modulation of intestinal stem cell niche morphogens alone can exert a neoplasia-relevant selective pressure on normal colonic epithelium. Using adult stem cell-derived murine colonic epithelial organoids (colonoids), we employed a strategy of sustained withdrawal of EGF and EGFR inhibition to select for and expand survivors. EGFR-signaling-independent (iEGFR) colonoids emerged over rounds of selection and expansion. Colonoids derived from a mouse model of chronic mucosal injury showed an enhanced ability to adapt to EGFR inhibition. Whole-exome and transcriptomic analyses of iEGFR colonoids demonstrated acquisition of deleterious mutations and altered expression of genes implicated in EGF signaling, pyroptosis, and CRC. iEGFR colonoids acquired dysplasia-associated cytomorphologic changes, an increased proliferative rate, and the ability to survive independently of other required niche factors. These changes were accompanied by emergence of aneuploidy and chromosomal instability; further, the observed mitotic segregation errors were significantly associated with loss of interkinetic nuclear migration, a fundamental and dynamic process underlying intestinal epithelial homeostasis. This study provides key evidence that chromosomal instability and other phenotypes associated with neoplasia can be induced *ex vivo* via adaptation to EGF withdrawal in normal and stably euploid colonic epithelium, without introducing cancer-associated driver mutations. In addition, prior mucosal injury accelerates this evolutionary process.

**Key definitions:** <u>Colonoids</u>: adult stem cell-derived colonic epithelial organoids

<u>iEGFR</u>: *in vitro* selective conditions devoid of EGF (epidermal growth factor) and including an EGFR (EGF receptor) inhibitor ^1^

<u>iEGFR colonoids</u>: colonoids tolerant to iEGFR culture conditions with growth and survival similar to unselected passage-matched controls

<u>INM</u>: Interkinetic nuclear migration

## INTRODUCTION

While much is known about the molecular features of CRC and their adenomatous precursors, it remains a mystery how neoplasia arises from normal epithelium^2,3^. The colonic epithelial crypt is a test-tube shaped unit comprised of Lgr5^+^ stem cells at its base, with its differentiation axis determined by epithelial and stromal microenvironment-derived gradients of niche growth factors^4–6^. How perturbations to normal niche growth factor homeostasis may act to promote or constrain initiation of epithelial neoplasia remain largely unexplored.

Advances in intestinal organoid culture have provided profound insights into the niche signaling pathways required for maintenance of epithelial homeostasis, including the EGFR/MAPK, Wnt, Notch, PI3K, and TGF-β pathways^7^. Intriguingly, these same pathways are recurrently altered in colorectal cancer (CRC), which is in turn characterized by epithelial architectural complexity, niche remodeling, and progressive loss of dependence on key niche factors^2,3,8,9^. Organoid cultures derived from adenomatous precursors and CRC demonstrate heterogeneous patterns of niche factor-independent growth reflective of underlying molecular changes^3,8^. For example, unlike normal epithelial cells, the vast majority of adenoma and CRC cells grow independently of Wnt and R-spondin *ex vivo*, underscoring the fact that *APC* mutation is a common first hit^3^. Further, multiple groups have leveraged the intestinal organoid model to reconstitute the adenoma-carcinoma sequence *in vitro*, harnessing selective strategies to identify successfully edited clones; for example, *KRAS* or *PIK3CA* mutant organoids survive in EGF-deficient conditions^10–12^.

The positioning of cells in the intestinal crypt dictates cell fate^13^. Interkinetic nuclear migration (INM) has recently been shown to contribute to the dynamics of cell positioning in the intestinal crypt^14^. INM is a homeostatic mitotic mechanism in intestinal epithelium by which basally located nuclei migrate to the apical aspect of the cell for mitosis, then return to a basal cytoplasmic location after separation of mitotic sisters^14^. Interestingly, loss of INM in *Apc* mutant murine intestinal organoids resulted in placement of mitotic sisters directly adjacent to one another, rather than the physically separated mitotic sisters seen in wild-type mitoses with intact INM^14^. Thus, in addition to altered niche growth factor homeostasis, biophysical factors related to mitotic dynamics, cell geometry, and/or microenvironmental stiffness may also directly contribute to clonal expansion of crypt cell populations to promote neoplasia. In the human colon, cycles of mucosal injury and repair (for example, in inflammatory bowel disease, or IBD) can transiently or permanently alter the biophysical properties, cell populations, and growth factors present in the mucosal microenvironment^15^. Although the mechanisms are not fully elucidated, such chronic inflammatory insults lead to increased risk for CRC and other epithelial cancers^16,17^.

Here we tested the hypothesis that disturbances to the mucosal microenvironment alone have capacity to lead to epithelial-autonomous molecular changes promoting cancer. As the feasibility of short-term EGF withdrawal in organoid culture has been demonstrated^1^, EGF is a critical intestinal stem cell niche factor^7,18–20^, and EGF signaling is indispensable for normal intestinal stem cell survival and propagation *in vitro*^3,7^, we focused our selection experiments on the evolution of EGFR-signaling-independent growth. Indeed, our data show that long-term withdrawal of EGFR signaling alone results in a molecularly distinct and sustained adaptive epithelial phenotype.

## METHODS

### Mouse Strains Used to Derive Colonoid Lines

Primary colonoid cultures used in this study were derived from C57BL/6J mice (directly received from Jackson Laboratory). All animal experiments were implemented in accordance with an animal protocol approved by the Johns Hopkins University Animal Care and Use Committee (Protocol MO18M85) and ARRIVE (Animal Research: Reporting of *In Vivo* Experiments) guidelines. All the mice were housed in a specific pathogen-free (Helicobacter negative) environment. The DSS chronic colitis mouse model was established as has been described previously with minor modifications^21–23^. In short, male mice 6-8 weeks old were treated with 4 rounds of DSS challenge, each consisting of 5 days of DSS in drinking water (2%, 40 kDa) followed by 7 days of recovery (Alfa Aesar #J63606).

### Colonoid Culture

We derived colonoids from normal wildtype C57BL/6 mice (referred to hereafter as control colonoids). Colonoids were derived from the distal 2.5 cm of grossly normal appearing C57BL/6J mouse colons (females 26 weeks old for control and *Apc*^mut^ colonoid lines). Absence of deleterious coding mutations was confirmed by whole-exome sequencing (data not shown). Notably, mouse colons lack Paneth cells, a potential source of EGF^24^. Colonic crypts were isolated and cultured as described previously^7,10^. Colonoids were plated within Matrigel (Corning #356231). Basic culture medium was composed of advanced Dulbecco’s modified Eagle’s medium/F12 (Gibco™) supplemented with penicillin/streptomycin, 10 mM HEPES (Gibco™ #15630080), GlutaMAX supplement (Gibco™ #35050061), B27™ Supplement (Gibco™ #17504044) and 1 mM N-acetylcysteine (Sigma-Aldrich #A9165). WENR medium was made of basic culture medium (20% final volume), Wnt3a-conditioned media (50% final volume with 5% final FBS concentration, L Wnt-3A ATCC® CRL-2647), and R-Spondin1-conditioned media (20% final volume), Noggin-conditioned media (10% final volume) and EGF (50 ng/mL). WNR medium had EGF omitted. Colonoids were maintained and propagated in culture as described previously^7^.

### Colonoid transfection and genome editing

The colonoid lipofection and CRISPR/Cas9 genome editing protocol was followed as described previously^25,26^. The sgRNA sequence targeting *Apc* can be found in Supplementary Figure 3A. As described previously, single colonoid survivors in Wnt/R-spondin-deficient media were manually picked and clonally expanded under the same selective conditions. The presence of biallelic truncating mutations at the expected site was confirmed by Topo cloning and whole-exome sequencing (Supplementary Figure 3B). Off-target coding mutations were not detected (data not shown).

### Derivation of iEGFR-tolerant organoids

Three days after plating in Matrigel, passage-matched colonoids were switched from WENR to iEGFR media (WNR with 5 μM Gefitinib; Santa Cruz #sc-202166). This concentration of gefitinib was required to kill >90% of normal colonoids at 7 days (data not shown) and was previously used to achieve iEGFR intestinal organoid culture conditions^1^. Fresh media with the drug was added every other day. Survivors were collected after 7 days and allowed to expand in WNR media before re-challenging in iEGFR selection for another 7 days. These cycles of selection and expansion were repeated until the survival rate plateaued (iEGFR-tolerant colonoids). All control colonoids were treated with similar concentration and volume of the compound dissolvent, dimethyl sulfoxide (DMSO, Corning® #25-950-CQC). Control colonoids were maintained in WENR media, and iEGFR-tolerant colonoids were maintained in WNR media long-term. Brightfield images of each cycle were captured on day 0 and day 7 using a Zeiss Microscope (Carl Zeiss Axiovert 40 C) with a 4x objective and the iDu Optics LabCam Microscope Adapter for iPhone8+ (iDu Optics). Quantification of survival rate was carried out manually. At the beginning of each cycle, the total number of colonoids in both control and treatment groups were counted under the microscope with a cell counter based on visual inspection (see images in Supplementary Figure 3D). At the end of each selection cycle (7 days), the total number of live colonoids in each well of both groups was counted. Survival rate was calculated as the total number of live colonoids post-treatment at day 7 to total number of colonoids pre-treatment at day 0 in each well. The relative survival rate was generated by comparing survival to untreated controls.

### Histology

Whole colonoids were collected by gently dissolving Matrigel in ice-cold PBS (pH 7.4), and subsequently fixed for 30 min at room temperature in 4% paraformaldehyde (16% PFA, Pierce™ #28906). Colonoids were then washed with PBS (pH 7.4) at room temperature. Colonoids were pelleted and transferred to the top of 2% solidified agarose gel in a 0.5 mL microfuge tube (Sigma-Aldrich #A9539). After aspiration of PBS, another 100uL of warm agarose gel was added to the colonoids. After gel solidification, the entire microcentrifuge tube was placed into a 15 mL conical containing 10 mL buffered 10% formalin (Sigma-Aldrich #HT501128) overnight. The bottom of the microcentrifuge tube was carefully removed with a razor blade and the colonoid block was transferred into a tissue cassette and submitted for paraffin embedding. 4 μm thick sections were stained with hematoxylin and eosin (performed by the Johns Hopkins Oncology Tissue Services Core). Photomicrographs of colonoids and deidentified human tissue samples (in accordance with the Johns Hopkins University School of Medicine Institutional Review Board, IRB00273344) were taken using an Olympus BX46 upright microscope and Teledyne Lumenera Infinity Analyze software.

### Metaphase spreads

Colonoids were treated with 100 μM colcemid (Gibco™ #15212012) for 4 hours and dissociated with 800 μL of TrypLE (Gibco™ #12604013) and Accutase (Invitrogen™ #00-4555-56) (1:1 ratio) for 10–15 min at 37 °C. After washout of TrypLE and accutase with advanced DMEM/F12 medium (Gibco™ #12634010) containing HEPES buffer (Gibco™ #15630080, 1 mM), penicillin/streptomycin (Gibco™ #15140122, 1%), GlutaMax (Gibco™ #35050061, 0.2 mM), cells were treated with pre-warmed KCl (0.56%) for 15 min at room temperature. Subsequently, 120 μl of fixative solution (methanol:acetic acid; 3:1) were added before centrifuging. After centrifugation, 10 ml fixative solution were slowly added before incubation at 4°C overnight. Fixed cells were dropped onto a glass microscope slide using a 20-μl pipette, air dried, and heat-dried (65°C) for 60 min. Slides were then incubated for 1 hour at 37 °C in propidium iodide (PI)/RNase staining buffer and rinsed with ddH_2_O. Slides were mounted with Vectashield containing DAPI (Vector Labs #H-1000) and analyzed on a Nikon laser microscope (×60 Super-Plan APO oil 1.4 NA objective). Control colonoids were assayed at passages 8 and 15; iEGFR colonoids were assayed at passage 40 (low, L) and passage 66 (high, H); *Apc*^mut^ colonoids were assayed at passages 28 and 30; DSS control colonoids were assayed at passages 5, 15, and 35; and DSS iEGFR colonoids were assayed at passages 15 and 35. Results were similar across passage numbers and combined per group, with the exception of iEGFR colonoids (as noted in Figure 4A). Chromosomes from each spread were manually counted in a blinded manner using Fiji/ImageJ.

### 3D immunostaining and clearing of organoids

Whole colonoids were collected by gently dissolving Matrigel in ice-cold PBS and fixed for 30 min at room temperature in 4% paraformaldehyde (PFA, Sigma). Colonoids were then transferred to organoid washing buffer (PBS containing 0.1% Triton X-100 and 0.2% BSA), then distributed into a 24-well plate. For immunofluorescent staining, colonoids were permeabilized and blocked in PBS containing 0.5% Triton X-100 and 1% BSA (Sigma) for 1 hour at room temperature, then incubated in blocking buffer containing primary antibody overnight at 4°C. Primary antibodies used were Chromogranin A (Santa Cruz #Sc-1488) and phospho-histone H2A.X (Ser139; Cell Signaling Tech #2577). Colonoids were incubated with corresponding secondary antibody Alexa 488 anti-mouse IgG (Invitrogen™ #A11029), in blocking buffer for overnight at 4°C, with 1ug/ul DAPI added for the final 15 minutes of incubation. Colonoids were washed 4-5 times (2 hours each), then cleared in fructose-glycerol clearing buffer (60% (vol/vol) glycerol and 2.5 M fructose) for 15 mins before imaging on a Zeiss LSM 780 confocal microscope^27^. Image analysis was performed using Zen and Fiji/ImageJ software.

### EdU incorporation assay

Colonoids were plated in an 8-well Chamber Coverglass (Nunc™ Lab-Tek™, Cat#155411). 4-5 days after plating, EdU (10 μM) was added to fresh medium for 6 hours. Colonoids were then fixed with warm 4% PFA for 10 mins at 37°C, then rinsed once with room temperature PBS. Blocking and permeabilization buffer (PBS containing 1% BSA and 0.5 % Triton X-100) was added for 2 hours at room temperature. EdU detection reagents were then added for 2 hours at room temperature in the dark (Click-iT™ Assay Kit, Sigma-Aldrich #C10337). Nutlin was used as a positive control (Selleckchem #S7101). Images were captured with the Zeiss LSM780 confocal microscope (40x/1.4 NA objective). Image analysis was performed using Zen, Fiji/ImageJ and Imaris software.

### Four-dimensional colonoid imaging and image analysis

#### Lentivirus production

The plasmids used were pMD2.G (Addgene plasmid RRID# 12259), psPAX2 (Addgene plasmid RRID#12260), and pLV-H2B-Neon-ires-Puro (kindly gifted by the Hugo J.G. Snippert and Geert J.P.L. Kops laboratories of the University Medical Center Utrecht). To make lentivirus particles, HEK 293FT cells were co-transfected with the lentiviral transfer plasmid, packaging plasmid, and envelope plasmid. Media containing lentivirus was collected 24 and 48 hours after transfection. Lentivirus was concentrated using a centrifugal filter (Amicon Ultra-15, 100,000 NMWL). The lentiviral titer was determined by qPCR (abm qPCR Lentivirus Titration Kit, cat. # LV900). Viral titers used in this study ranged from 1×10^8^-1×10^9^ IU/ml.

#### Lentiviral infection of colonoids

To visualize mitoses, colonoids were infected with lentivirus encoding mNeon-tagged histone 2B and a puromycin resistance cassette described above. The protocol was performed as described previously with minor modifications^28^. Briefly, colonoids ~100 μm in diameter were transferred to a 15 ml tube and pelleted (1000 rpm for 5 minutes) before single cell dissociation (600-800μL TrypLE, 37°C). Pelleted single cells were resuspended in 1 ml of prewarmed infection medium, consisting of 500μL concentrated virus, 500 μl WENR (control colonoids), WNR (iEGFR colonoids), or Wnt/R-spondin deficient media (*Apc*^mut^ colonoids), 8 μg/ml Polybrene (Sigma-Aldrich #TR-1003), and 10 μM Rock inhibitor Y-27632 (Sigma-Aldrich #Y0503), then centrifuged at 100 rpm for 1h at room temperature. Colonoids were then transferred to the cell incubator (37°C, 5% CO2) for 5–6 hours and gently remixed every hour prior to replating with fresh media as indicated. Approximately 2-3 days after infection, the expression of transduced fluorescence protein was observed and puromycin selection (1 μg/mL) was initiated. Puromycin was increased to 5ug/mL once colonoid size reached more than 100 μm.

#### Four-dimensional colonoid imaging

After two passages of puromycin selection, colonoids were dissociated using TrypLE and replated in an 8-well glass-bottom chamber slide (Nunc™ Lab-Tek™, Cat#155411). Three to four days later, the chamber was mounted on a confocal laser-scanning microscope (LSM 780), which was continuously held at 37 °C with 5.0% CO_2_. H2B-Neon-positive organoids were imaged in xyzt mode for 16–18h at 37°C at 3 min intervals using a 40x water-immersion objective (NA 1.1). Eight to ten H2B-mNeon-expressing colonoids were imaged simultaneously using minimal amounts of 488 nm laser excitation. In total, 14-16 z-sections at 2-μm intervals were imaged per colonoid.

#### Imaging analysis

To analyze mitoses, raw image Z-stacks were converted to depth color-coded maximum projections with using a custom macro modified from the ImageJ/Fiji software plugin “Temporal-Color Code”^29^. The macro attributes a color code to each z-layer, facilitating visual discrimination of cells overlapping in XY as described previously^10^. Data sets were converted into manageable and maximally informative videos, combining z-projection, depth color-coding and merging with transmitted light images (Supplementary Videos 1–6). Mitoses were blinded and scored, judged and counted manually by both T.C. and T.C.L. For analysis of interkinetic nuclear migration, Fiji/ImageJ was used to measure the pixel distance the basal aspect of a nucleus moved prior to mitotic entry and nuclear envelope breakdown. Any distance moved was categorized as intact interkinetic nuclear migration. No measurable movement was categorized as loss of interkinetic nuclear migration.

### Quantitative RT-PCR based mouse karyotyping

SYBR Green qPCR assays were designed and validated for every mouse chromosome based on GRCm38/mm10 genome assembly (primer sequences listed in Table S1). qPCR reactions were set up in triplicate in a 384-well plate and run on the CFX384 Touch Real-Time PCR Detection System (Bio-Rad). Each reaction contained 5μL of PerfeCTa SYBR Green FastMix (Quantabio, catalog number 95073-05K), 2.5μL of forward and reverse primer mix at 2μM, 0.5μL of purified genomic DNA at 1ng/μL and 2μL of nuclease-free water. A standard cycling protocol was followed as provided with the SYBR Green reagent. Ct values were acquired with CFX Manager Software (Bio-Rad) and the relative chromosome copy numbers were calculated using a modified ΔΔCt method as described previously^30^.

### RNA sequencing

Total RNA was isolated from pelleted colonoids in using Trizol (Invitrogen™ #15596026) according to the manufacturer’s instructions and purified using the Purelink RNA Mini Kit (Invitrogen™ #12183018A). RNA-sequencing data were generated by Novogene. cDNA libraries were sequenced on an Illumina NextSeq500 using 75-bp paired-end sequencing. Clean reads were mapped to UCSC GRCm38 reference genome using STAR v2.5 software^31^ and raw counts were assigned to Ensembl genes using featureCounts (subread v2.0.0 aligner command line tool)^32^. Differential gene analysis was performed using DESeq2 v1.28.1 following regularized logarithm transformation of raw count data^33^. Gene set enrichment analysis (GSEA) was performed using the gost function of the R package gprofiler2 v0.1.9^34^. Genes were considered differentially expressed and included in the GSEA if they had a *p*-value <3.3e-7 (Bonferroni adjusted) and an absolute log_2_-transformed fold change >2. Statistical analysis and plotting were performed using R software (version 3.4.0). Statistical significance was assessed at α=0.05. The significance level for differential gene analysis was adjusted using the Bonferroni approach while also accounting for multiple comparisons across experimental conditions (threshold *p*=3.3e-7). All data are presented as mean ± SEM. Analysis of two samples was performed with unpaired two-tailed student t-test for equal variance, or t-test with Welch’s correction for heterogeneity of variance.

### Whole Exome sequencing

DNA was extracted from pelleted colonoids using the Purelink Genomic DNA Kit (Invitrogen™ #K1820-01) according to the manufacturer’s instructions. Whole exome sequencing data were generated by BGI. In short, fragmented gDNA was subjected to adapter ligation, amplification, and exome array hybridization. Captured products were circularized and DNA nanoballs were produced using rolling circle amplification prior to loading onto the BGISEQ sequencing platform. Mean sequencing depth on target regions was 117.58x, and 98.69% of targeted bases had at least 10x coverage. Paired-end reads were mapped to UCSC GRCm38 and aligned using Burrows-Wheeler Aligner (BWA) software. The Genome Analysis Toolkit (GATK) was used for variant calling and the SnpEff tool was used for variant annotation. Variants of interest were filtered based on >10x depth of coverage, predicted high functional impact (MutDB), and visual inspection using Integrative Genomics Viewer (IGV).

### Quantitative RT-PCR

cDNA was isolated using SuperScript™ III Reverse Transcriptase (Invitrogen # 108080) following the manufacturer’s protocol. A dilution series of cDNA was used to validate the primer pairs, and both melting curve analysis and agarose gel electrophoresis were performed to check for specificity of primers (data not shown). Quantitative PCR was performed using the SYBR green Select Master Mix (Thermo Fisher #4472908) following the manufacturer’s protocol. Each sample was done in triplicate in a total reaction volume of 10-μl containing 0.5-μl of 1:8 diluted cDNA (validated dilution with dynamic range of amplification) and 5nM primer mix using the CFX384 QPCR machine (Biorad). The list of primers used are listed in Supplementary Table 3. Delta (Cq) was calculated by subtracting the mean Cq for every tested gene to those of the internal control genes (*Hprt* and *ActB*). Log fold change was calculated by subtracting Delta(Cq) of iEGFR samples from those of controls.

## RESULTS

### Normal murine colonoids can achieve sustained EGFR-independent growth in long-term culture

We tested our hypothesis that changes in the availability of niche factors can select for a cancer phenotype in normal wild-type colonic epithelium using colon-derived organoids (colonoids). To select for EGFR-independent growth *in vitro*, we cultured normal colonoids from wild-type mice (“control” colonoids) in EGF-depleted (WNR) medium with the EGFR-specific inhibitor Gefitinib; we refer to these culture conditions as iEGFR. The EGFR inhibitor was used to address the possibility of autocrine/paracrine production of EGF by the cultured colonoids or the presence of exogenous EGF in the 5% final concentration of fetal bovine serum in EGF-depleted medium. Over 7 days, iEGFR selection resulted in the death of most colonoids. Rare survivor colonoids appeared smaller and lacked budding compared to control colonoids, suggesting that they were mostly quiescent^1^. These survivors were recovered and expanded in WNR media, but not in the presence of Gefitinib. We continued to re-challenge the expanded survivors with additional 7-day cycles of iEGFR selection. Increasing numbers of survivors were recovered with each re-challenge cycle (Figure 1A), and approximately half of colonoids survived after 3 cycles of selection. A total of 5 cycles were required to achieve complete EGFR-independent growth (iEGFR colonoids), with survival rate similar to unchallenged colonoids in EGF-replete media (Figure 1A). Notably, we also observed a transient enrichment of cells with enteroendocrine differentiation during iEGFR selection as reported previously (Supplementary Figures 1A-B)^1^. We continuously propagated iEGFR colonoids for 8 months in WNR medium.

**Figure 1.**
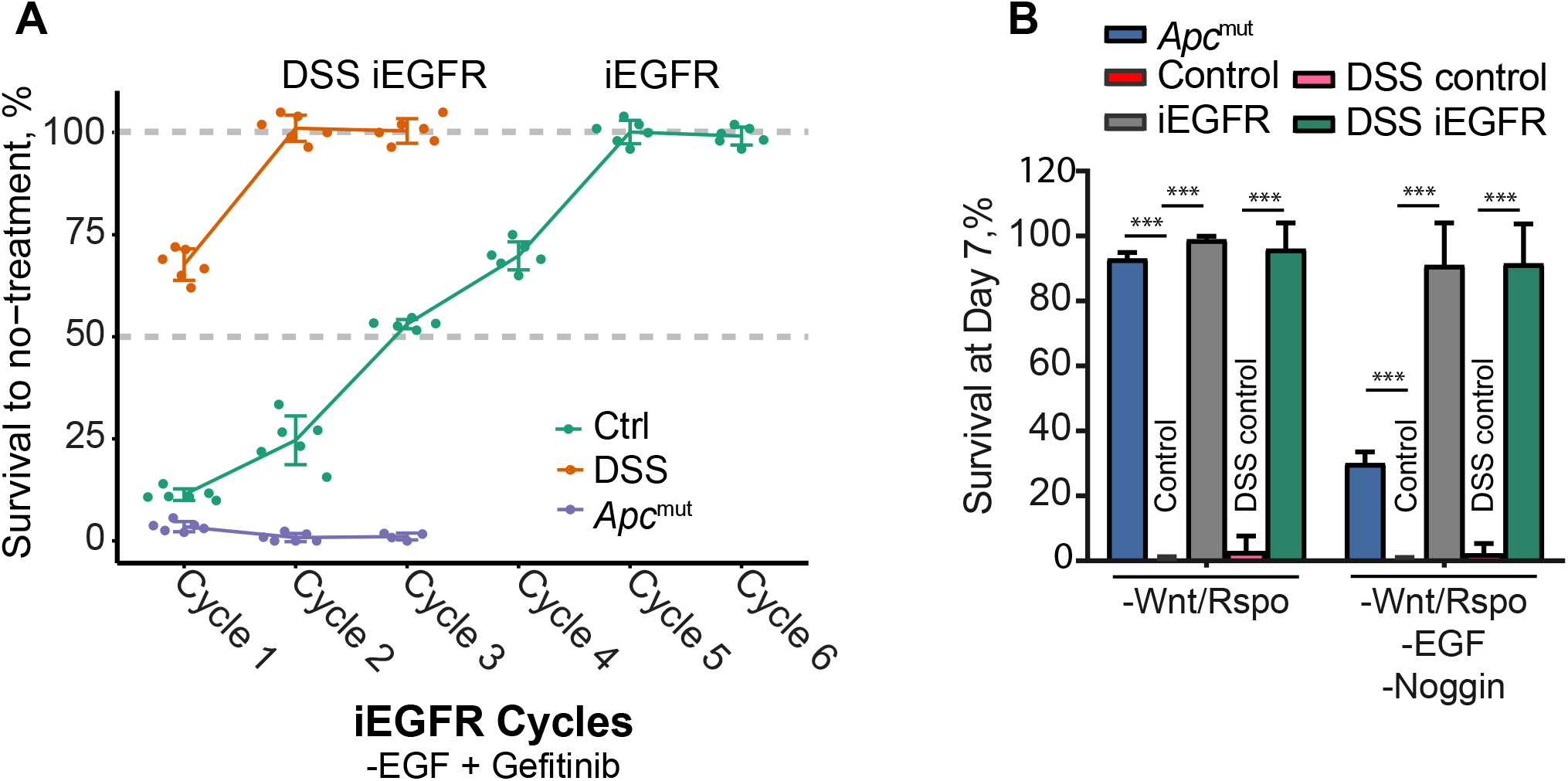
Colonoids adapt to culture conditions devoid of critical niche factors. **A)** Survival rate depicting viability of colonoids at the end of each selection cycle (7 days) as a percentage of colonoid growth in control media (n= 6 biological replicates). **B)** Survival rate of colonoid lines in other selective media after 7 days (n=3 biological replicates), **** p < 0.0001, 2-tailed non-paired student t-test.

To test whether iEGFR tolerance was reversible after relaxing selective conditions, we returned iEGFR colonoids to EGF-replete medium (WENR) for 3 weeks. Surprisingly, approximately 70% of iEGFR colonoids died each week in the presence of EGF compared to those maintained in the WNR medium. Moreover, removal of EGF from the culture reverted to the growth of iEGFR colonoid baseline of near-100% survival (Supplemental Figure 1C). Taken together, our data demonstrate the feasibility of evolving and propagating growth factor-independent colonic epithelium. They also show that the resulting phenotype is a stable trait that does not require persistent selection once acquired with an acquired and related vulnerability.

### Prior epithelial injury facilitates adaptation to iEGFR selection

We tested whether chronic injury and repair can influence adaptability to EGF deprivation using colonoids generated from a mouse model of chronic chemical colitis (DSS, dextran sodium sulfate). As previously described^35^, these mice showed cardinal signs of colitis as manifested by a significant reduction in the ratio between body weight to colon length (Supplementary Figure 2A) and histologic features (Supplementary Figure 2B). Colonoids derived from DSS-treated mice more readily adapted to iEGFR selection compared to controls, with approximately 60% of colonoids surviving the first 7 day cycle of selection compared to approximately 10% of control colonoids (Figure 1A). In addition, DSS colonoids reached a survival plateau after only 2 cycles of iEGFR selection (Figure 1B), compared to 5 cycles for control colonoids. These data indicate that prior exposure to cycles of mucosal injury *in vivo* primed colonic epithelium for adaptation to iEGFR selection.

### iEGFR colonoids acquire tolerance to deprivation of other niche factors

To test whether iEGFR colonoids more readily acquire additional niche factor independence, we challenged them in medium lacking Wnt/R-spondin, as well as in a base medium that additionally lacks Noggin. The majority of iEGFR colonoid lines survived this selective challenge after a week (Figure 1B, Supplementary Figure 3D). On the other hand, the majority of control colonoids did not survive either condition. These data demonstrate that iEGFR colonoids acquired the capacity to tolerate additional niche-relevant selective pressures.

### *Apc* mutant colonoids are more vulnerable to EGF-deficient conditions

Mathematical modeling of CRC carcinogenesis suggests that *APC* mutations may accelerate the acquisition of subsequent molecular alterations^36^. If iEGFR adaptation relies upon *de novo* oncogenic mutation, *Apc* loss should confer an adaptive advantage. To test this hypothesis, we introduced biallelic truncating mutations in *Apc* via CRISPR/Cas9 genome editing to a control mouse colonoid line (Supplementary Figure 3A-B, confirmed by whole exome sequencing). As previously described, *Apc* mutant (*Apc*^mut^) colonoids grew independently of Wnt/R-spondin-containing medium and adopted spheroid morphology (Figure 1B and Supplemental Figure 3C)^25^. Surprisingly, when *Apc*^mut^ colonoids were subjected to cycles of iEGFR selection, unlike wild type colonoids, *Apc*^mut^ colonoids could not adapt to EGF deprivation (Figure 1A). These data suggest that *Apc* loss greatly enhances the sensitivity of colonoids to EGFR deprivation.

### iEGFR colonoids acquire somatic mutations and transcriptional reprogramming

To determine whether mutations associated with EGF-independent growth may have contributed to the iEGFR phenotype, we performed whole exome sequencing on iEGFR colonoids and controls. First, we looked at mutations in the EGFR signaling pathway that are frequently observed in CRC. No mutations or indels in *Egfr, Kras,* or *Pik3ca* were detected in iEGFR colonoids. However, we detected a coding mutation with predicted high functional impact in *Wnk2,* a negative regulator of EGF-induced activation of ERK/MAPK signaling^37^. iEGFR colonoids also showed predicted deleterious mutations in *Btk,* known to have a role in negatively regulating Wnt-β-catenin signaling^38^, *Treml2, and Olfr1255* (Table 1). Mining publicly available data reveals that these genes are altered at very low frequency (<1%) in human CRC, with the exception of *Olfr1255* (Table 1). DSS iEGFR colonoids were enriched for a mutation in *Ninein* (Table 1), a gene involved in centrosomal biology and mitotic fidelity^39^. DSS colonoids did not accumulate coding mutations compared to control colonoids.

**Table 1.**
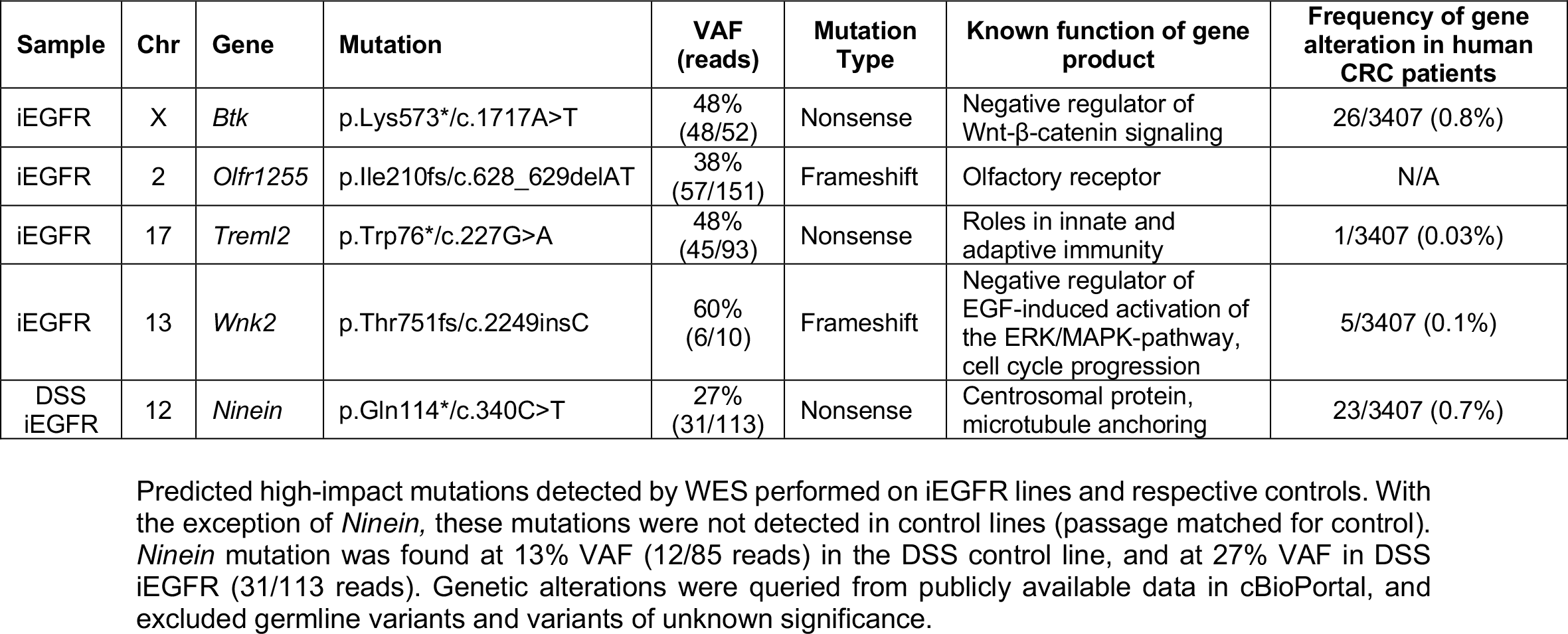
Summary of mutations identified in iEGFR colonoids by WES.

We also performed RNA-sequencing to explore the molecular changes in iEGFR, DSS iEGFR lines, and passage-matched controls. Principal component analysis established that replicate samples clustered together with high reproducibility (Figure 2A). We found a total of 547 differentially expressed genes in both types of iEGFR colonoids (absolute log_2_ fold change >2, Bonferroni-adjusted p-value < 0.05, Figure 2B, Supplementary Table 1, Supplementary Figure 4). Gene set enrichment analysis of the upregulated overlapping genes (48) showed significant enrichment for genes associated with amine transmembrane transporter activity, pyroptosis, and phosphatidylinositol-4-phosphate binding pathways, while the overlapping downregulated genes (22) were involved in endocytosis and cell junction assembly (Figure 2C). A subset of significantly differentially expressed genes was further validated using quantitative reverse transcription polymerase chain reaction (qRT-PCR), including those with roles in the EGF pathway, pyroptosis, and CRC carcinogenesis, such as *Igfpb7*, *Efemp1, Gasdmc2, and Mycn* (Figure 2D; all tested genes validated). Taken together, these data show that key neoplasia-relevant gene expression patterns emerge in colonic epithelial cells tolerant to EGF withdrawal.

**Figure 2.**
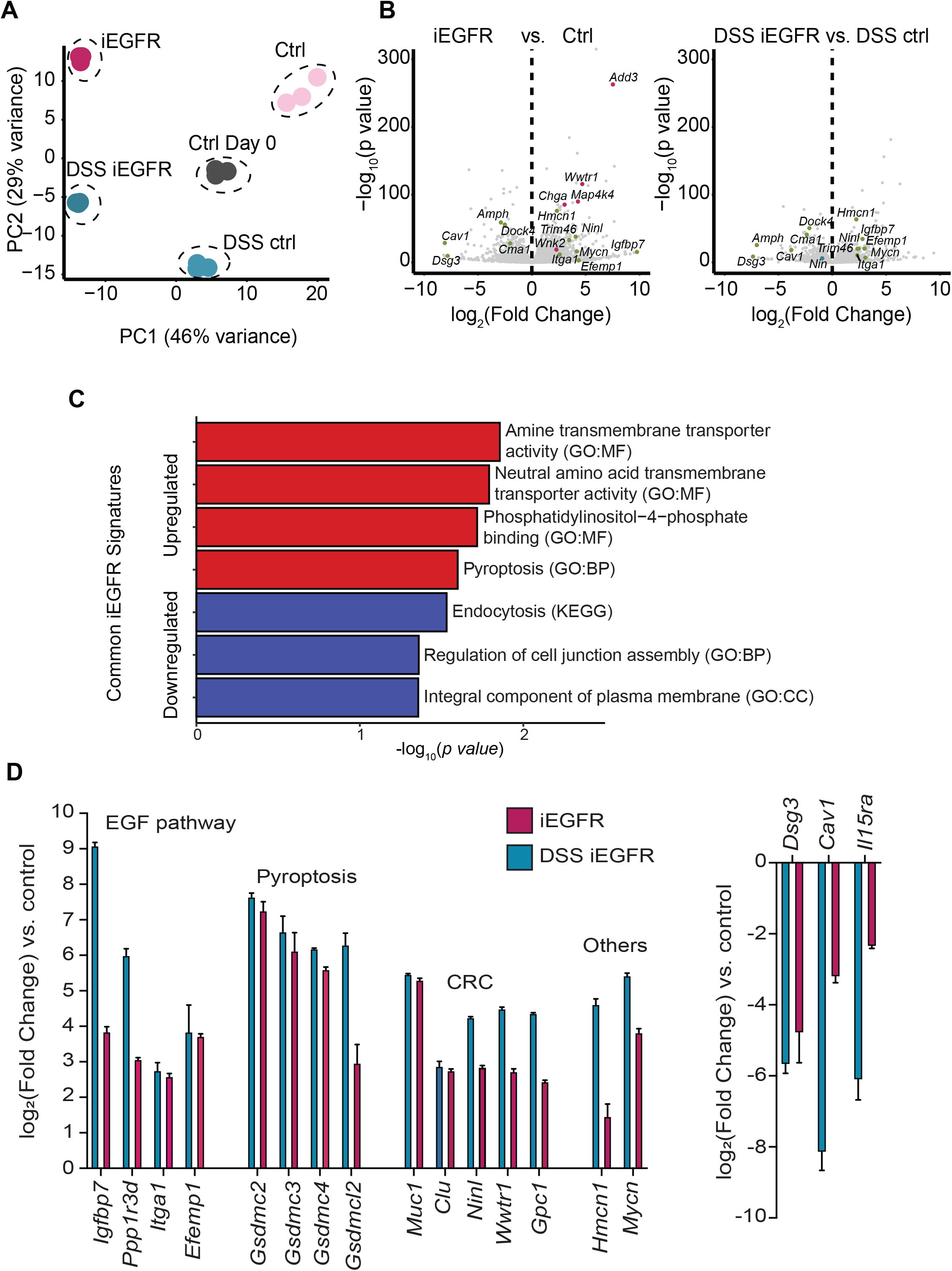
Long-term adaptation of colonoids to EGF-deficient conditions results in transcriptional changes. **A)** Samples analyzed by RNA-seq were plotted by principal component 1 (PC1) and principal component 2 (PC2) using raw count data following regularized logarithm transformation. Samples from the same experimental condition were grouped with the same colors. **B)** Volcano plots displaying log_2_-transformed fold change and −log_10_-transformed *p* value of genes assessed by RNA-seq in iEGFR vs. control colonoids and DSS iEGFR vs. DSS control colonoids. Selected differentially expressed genes are highlighted. Genes highlighted in green are differentially expressed in both iEGFR and DSS iEGFR compared to the respective control. Genes highlighted in red and blue are differentially expressed only in iEGFR or DSS iEGFR, respectively. **C)** Gene set enrichment analysis of overlapping upregulated and downregulated genes in both iEGFR and DSS iEGFR compared to the respective control. All enriched gene sets (*p* value < 0.05) are shown. **D)** Quantitative RT-PCR validation of select upregulated (left) and downregulated (right) genes detected by RNA-seq. Results are expressed as log_2_ fold change to control and DSS control (n=3).

### iEGFR colonoids exhibit morphologic changes and increased proliferation

Early epithelial neoplasia demonstrates characteristic morphological changes that have been routinely used by pathologists to diagnose dysplasia and cancer for over a century^40^. Hematoxylin-and-eosin-stained iEGFR colonoids show heterogeneous morphologic changes associated with dysplasia, including nuclear hyperchromasia, pseudostratification, and increased nuclear-to-cytoplasmic ratio (Figure 3A). DSS control colonoids showed features of reactive and regenerative epithelium, including more squamoid cells with brightly eosinophilic cytoplasm. A subset of DSS iEGFR colonoids strikingly showed loss of polarity and architectural complexity reminiscent of high-grade colitis-associated dysplasia seen in patients with IBD (Figure 3A).

**Figure 3.**
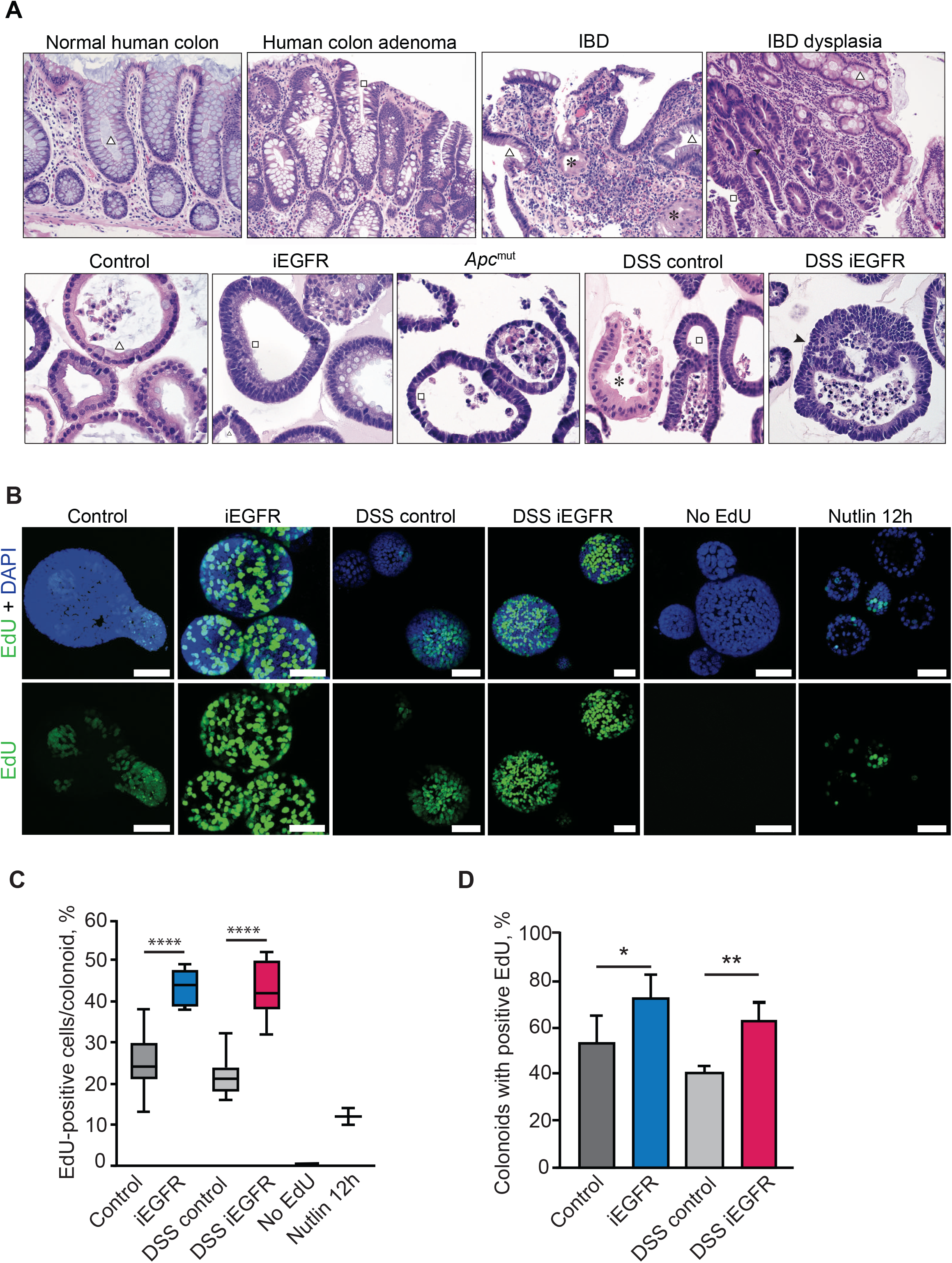
iEGFR colonoids show morphologic features of dysplasia and increased proliferation. **A)** Representative H&E stained human tissues (upper panels, 20X) and cultured colonoids (lower panels, 40X). Squares denote nuclear hyperchromasia and loss of nuclear polarity, arrowheads denote architectural complexity, asterisks denote squamous features, and the triangle denotes overall normal epithelial morphology. **B**) Representative confocal maximal Z-stacks images for colonoids stained with the thymidine analogue EdU (green) and the counterstain DAPI (blue). No-EdU and Nutlin served as negative and positive controls, respectively. Scale bars = 50 μm. n = 3 independent experiments. C) Box and whiskers plot for the percentage of EdU positive nuclei per EdU-positive colonoid. Transverse lines represent the median, boxes show 25^th^-75^th^ percentile and the whiskers represent the lowest and highest values within 1.5 times the interquartile range. **** *p* < 0.0001, \**p* ≤ 0.05; 2-tailed, non-paired student t-test. **D)** Bar plot for the percentage of colonoids with at least one EdU positive cell (n= 3 biological replicates). Error bars represent standard deviation. \*\*\*\**p* < 0.0001, \**p* ≤ 0.05; 2-tailed, non-paired student t-test.

Neoplasia is also associated with sustained proliferation^41^. Our RNA-sequencing analysis revealed that many genes associated with cellular proliferation were upregulated in iEGFR colonoids, including the proto-oncogene *Mycn* (Figure 2D, Supplementary Table 1). To further explore this, we used a short-pulse (6 hours) of the nucleotide analogue 5’-ethynyl-2’deoxyuridine (EdU) to analyze the proportion of cells in the S-phase of the cell cycle in iEGFR colonoids. The proportion of EdU+ colonoids and percent of EdU+ cells per colonoid were significantly higher in both iEGFR and DSS iEGFR colonoids compared to their controls (Figure 3B-3D). These data show that the iEGFR phenotype is characterized by histologic features of dysplasia and increased proliferation.

### iEGFR colonoids develop aneuploidy and chromosomal instability

Loss of genomic integrity is one of the hallmarks of cancer, and chromosomal instability (CIN, or ongoing aneuploidy) is observed in the majority of sporadic and IBD-associated CRC^2,41^. As aneuploidy fuels adaptation to selective pressures^30,42^, we hypothesized that this may play a role in acquisition of iEGFR tolerance. As the long-term genetic stability of adult stem cell derived intestinal organoid cultures has been established^43^, as expected, metaphase spreads of control organoids were mostly euploid (Figure 4A-B). In contrast, DSS-control colonoids were enriched for polyploidy (Figure 4A-B). Previous literature has implicated *APC* loss in promoting CIN^44^; we observed both polyploidy and aneuploidy in metaphase spreads of *Apc*^mut^ colonoids (Figure 4A).

**Figure 4.**
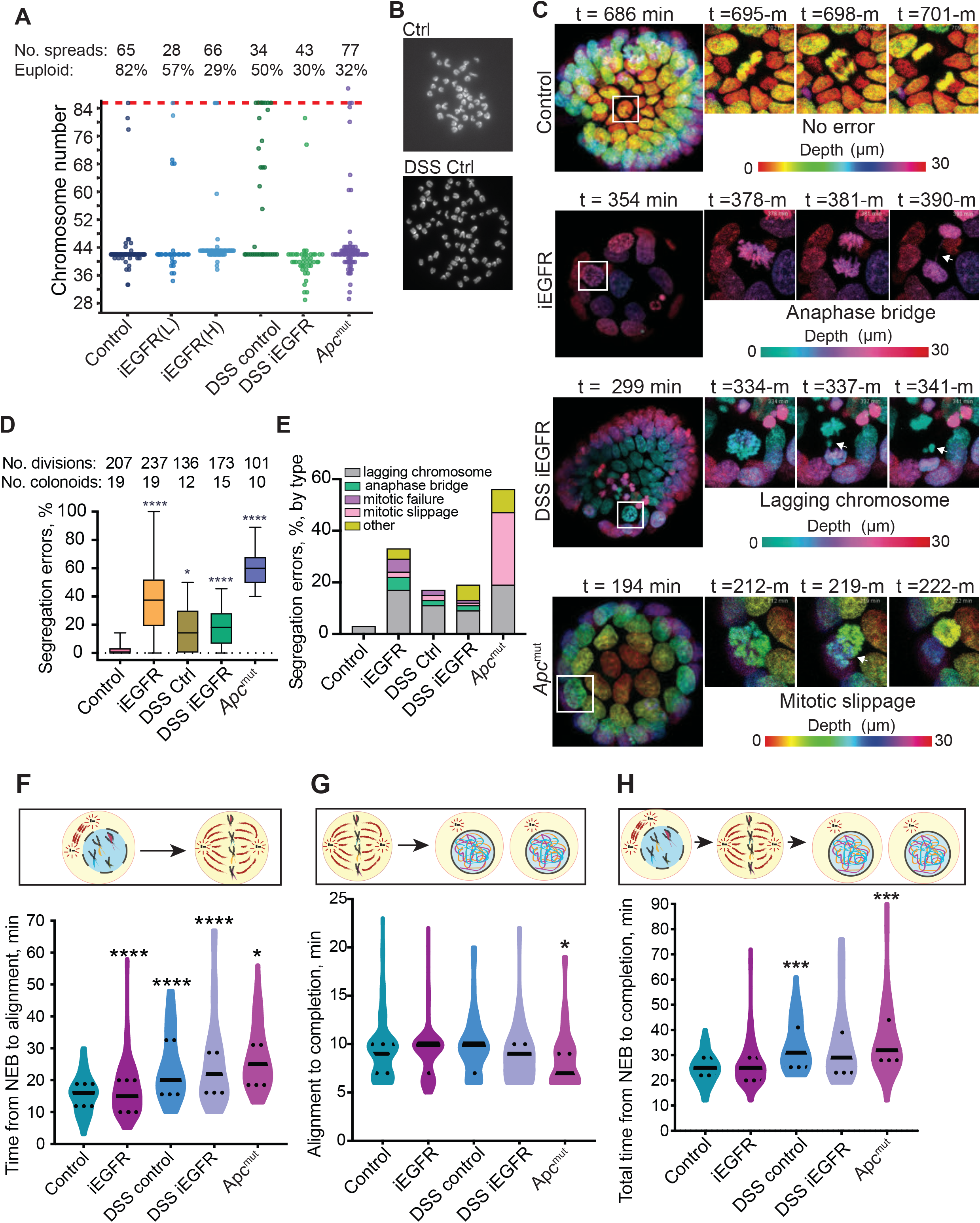
iEGFR colonoids are aneuploid and demonstrate chromosomal instability. **A)** Dot plot of the number of chromosomes in metaphase spreads. The number of counted spreads and the percentage of metaphase spreads with euploid chromosomes are shown at the top. The red line represents tetraploidy. iEGFR(H) and iEGFR(L) correspond to high passage number (‘H’ high, passage 66) and lower passage number (‘L’ low, passage 40), respectively. **B)** Representative images of metaphase spreads from control (euploid) and DSS control (tetraploid) colonoids. 60x. **C)** Representative color depth coded images of chromosome segregation errors as revealed by H2B-mNeon labeling of colonoids. Insets highlight mitoses in white boxes. White arrows indicate mitotic errors, corresponding to Supplementary Videos 1-6. n= 4 or 5 independent experiments. **D)** Box and whiskers plot of the percentage of segregation errors. Transverse lines represent the median, boxes show 25^th^-75^th^ percentile and the whiskers represent the lowest and highest values within 1.5 times the interquartile range. The number of divisions and colonoids analyzed are shown at the top. \*\*\*\**p* < 0.0001, \**p* ≤ 0.05; 2-tailed, non-paired student t-test. **E)** Bar plot of the percentage of different segregation errors in analyzed mitotic figures. Other types of errors include multipolar mitoses, mitotic failure, and fusion of nuclei. **F-H)** Illustrative cartoons and violin plots for time distribution of duration from nuclear envelope breakdown (NEB) to chromosome alignment (F), chromosome alignment to completion of mitosis (G), and total mitotic time (H). Transverse solid lines represent the median and the dotted lines border 25^th^ −75^th^ percentiles. \*\*\*\**p* < 0.0001, \**p* ≤ 0.05; 2-tailed, non-paired student t-test.

We observed that heterogenous aneuploidy arose during iEGFR selection, with an overall tendency for reduction in chromosomal number (subdiploid) (Figure 4A). Longer-term propagation of iEGFR colonoids (more than 25 additional passages) resulted in convergence onto a gain of one chromosome (Figure 4A, iEGFR ‘H’, or high passage)). Quantitative chromosome stoichiometry analysis via quantitative PCR revealed a complete loss of chromosome 13 at an earlier passage (~12 passages earlier than iEGFR ‘H’, Supplementary Figure 5). We hypothesized that aneuploidy could be due to increased DNA damage, but did not detect increased double stranded breaks as assessed by γH2AX staining in iEGFR colonoids relative to their corresponding controls (Supplemental Figure 6).

We next investigated the possibility that the heterogeneous aneuploidy was associated with ongoing CIN. The dynamic properties of mitosis were quantified via live-imaging of 3D colonoid cultures of H2B-mNeon expressing cells (Figure 4C-F, Supplementary Videos 1-6). The mean length of mitosis in control colonoids was ~30 minutes, with errors detected in only 3-5% of all mitotic events (Figure 4C-D, H). In contrast, iEGFR, *Apc*^mut^, DSS control, and DSS iEGFR colonoids showed a significantly elevated rate of erroneous mitoses relative to controls, ranging from approximately 20% (DSS lines) to 60% (*Apc*^mut^) (Figure 4D-E). In addition, the mean overall time of mitosis was significantly increased in DSS control, DSS iEGFR, and *Apc*^mut^ organoids compared to control colonoids (Figure 4H), mostly driven by increased length of nuclear envelope breakdown to chromosome alignment (Figure 4F-G).

Interestingly, altered mitotic timing was observed in DSS and *Apc*^mut^ lines but not in controls (Figure 4F-H), suggesting defects in the spindle assembly checkpoint in these colonoids.

Finally, live imaging revealed a normal pattern of INM in most control colonoids undergoing mitosis (Figure 5A), as previously described^14^. In contrast, this process was conspicuously absent in many iEGFR, DSS control, DSS iEGFR, and *Apc*^mut^ mitoses (Figure 5A). Further, INM loss was significantly associated with mitotic errors in our colonoid lines (Figure 5B-C).

**Figure 5.**
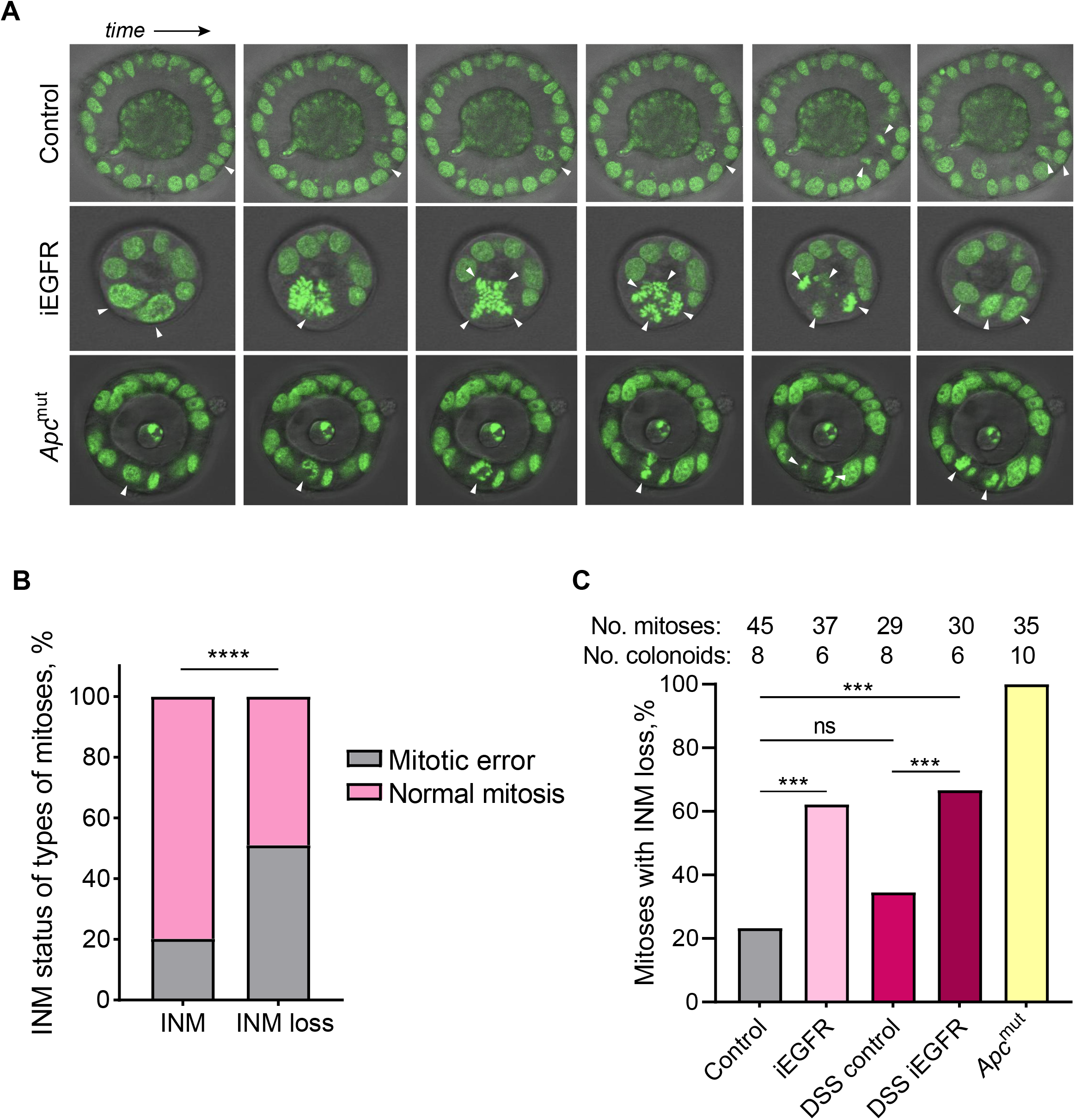
INM loss is frequent in iEGFR colonoids and significantly associated with mitotic errors. **A)** Sequential still images captured from representative individual mitoses (highlighted by white arrowheads) as revealed by H2B-mNeon labeling of control, iEGFR, and *Apc*^mut^ colonoids. **B)** Bar graph stratifying the presence of mitotic errors with the presence (n=79) or loss (n=98) of INM in all analyzed mitoses across colonoid lines. \*\*\*\**p* < 0.0001; 2-tailed Fisher’s exact test. **C)** Bar graph detailing the percentage of mitoses with INM loss in each colonoid line. The number of mitoses and colonoids evaluated per group are shown at top. \*\*\**p* < 0.001, \**p* ≤ 0.05; 2-tailed Fisher’s exact test.

Taken together, these data demonstrate that adaptation to long-term EGF withdrawal is associated with mitotic defects that result in chromosomal instability and aneuploidy. In addition, we associate loss of interkinetic nuclear migration with mitotic errors in colonic epithelium. Our data also show that prior chronic mucosal injury predisposes to epithelial chromosomal instability that persists *ex vivo*.

## DISCUSSION

Here, using long-term selective culture, we demonstrate that normal colonoids can adapt to withdrawal of the critical niche factor EGF, a process associated with cytomorphologic features of dysplasia, loss of INM, aneuploidy, CIN, somatic deleterious mutations, and transcriptional reprogramming. These data support a scenario in which epithelial-autonomous molecular changes known to be associated with neoplasia can arise during adaptation, the acquisition of which are accelerated by prior mucosal injury.

We discovered that iEGFR colonoids show aneuploidy and CIN (Figure 4) and are primed to adapt to other niche-relevant selective pressures (Figure 1B). Genomic copy number changes were recently shown to precede chronic inflammation-associated esophageal adenocarcinoma up to a decade prior to histologic evidence of transformation^45^, supporting an early initiating role for genomic instability. Recent work has demonstrated low levels of spontaneous cell fusions (as we observed in iEGFR lines) in cancer cell lines that led to increased phenotypic plasticity and accelerating adaptive potential^46^. Future work will determine whether CIN has a causal role in mediating iEGFR tolerance.

Carroll et al. demonstrated that INM is an important homeostatic mechanism involved in directing long-term cell positioning in the intestinal crypt^14^. While approximately one third of normal mitoses with intact INM led to separation of mitotic sisters, mitotic sisters always remained direct neighbors in the setting of *Apc* mutation, potentially contributing to clonal expansion of early adenomas. Our data validate their finding of INM loss with *Apc* mutation, which we further extend by associating INM loss with mitotic errors in colonoid lines (Figure 5). While our live imaging data precluded definitive evaluation of post-mitotic sister cell placement, it is possible that INM loss renders cells with mitotic errors more likely to undergo clonal expansion.

Inflammation and mucosal injury reprograms colonic epithelium to a regenerative/progenitor-like status^47–49^, and in other organ systems, stem cell lineage infidelity drives both wound repair (homeostatic) and cancer (pathogenic)^50,51^. Consistent with this literature, chronic DSS colonoids showed enhanced adaptive potential to iEGFR selective culture (Figure 1A). In addition, we found polyploidy in DSS colonoids, which has also been previously reported in the setting of wound repair (Figure 4A-B)^52,53^. Our RNA-sequencing analysis revealed that genes related to pyroptosis were significantly upregulated in iEGFR lines (Figure 2C-D). Pyroptosis, a caspase-dependent form of proinflammatory programmed cell death, has emerging roles in the tumor microenvironment and has been recently implicated in promoting colitis-associated cancer^54,55^. Recent investigation of somatic evolution in IBD colonic epithelium revealed clonal expansions of mutations in the IL-17 pathway which render epithelium resistant to the IL-17A-induced pro-apoptotic response^56,57^. Whether aneuploidy similarly confers resistance to pyroptosis-associated cell death is a future avenue of exploration.

Prior studies indicate that human adenoma-derived colonoids are uniformly dependent on EGF in culture, similar to normal colonoids^3^. We were surprised to find that, in contrast to wild-type control colonoids, *Apc*^mut^ colonoids were not able to overcome withdrawal of EGF (Figure 1A). While it is well known that *KRAS/PIK3CA* wild-type CRC is susceptible to EGFR inhibition^58^, our data suggest that this response may also be *APC* mutation-dependent.

Recent literature has demonstrated the presence of age-associated somatic mutations in normal non-dysplastic colonic epithelium across the lifespan with uncertain consequences^59–61^. Although rare patients with CRC harbor alterations in the somatically mutated genes we observed in iEGFR colonoids, whether these mutations act as drivers versus passengers in the adaptive iEGFR phenotype remains to be determined. We also acknowledge that bulk exome DNA sequencing may not detect rare mutations or mutations in regulatory elements that may contribute to the phenotype of our heterogeneous iEGFR colonoid lines.

Although further work is required to elucidate a potential role for perturbed niche homeostasis in human CRC initiation, our data support a potential role for microenvironmental selective pressures in promoting neoplastic transformation. Thus, increasing the granularity of our understanding of colon anatomic segment-specific mucosal microenvironments may reveal insights into the origins of distinct pathways of tumorigenesis (for example, the serrated versus adenomatous pathways of carcinogenesis in the proximal vs. distal colon, respectively).

In summary, we leveraged murine colonoids to demonstrate that sustained deprivation of niche-relevant growth factors alone can molecularly reprogram colonic epithelium. We anticipate that there are a spectrum of mechanisms epithelia can draw upon to adapt to such selective conditions. Tracking individual clones over time and extending our approaches to human and IBD-derived colonoid lines may determine whether adaptation mechanisms such as CIN are observed more broadly. Further, elucidating the mechanisms by which neoplasia-promoting epithelial phenotypes arise may reveal general vulnerabilities attractive for cancer prevention.

## SUPPLEMENTARY FIGURES, TABLES, AND VIDEOS

**Supplementary Figure 1. A)** Representative confocal fluorescence images (Z sections with maximum projection) of cleared colonoids labeled with chromogranin A (CHGA, green) and the counter stain DAPI (blue). **B)** Bar plot illustrating the number of chromogranin A positive cells per colonoid, evidence of enteroendocrine differentiation. Error bars represent standard deviation. *** *p* < 0.001. **C)** Bar plot demonstrating the survival rate of iEGFR colonoids after re-challenging with iEGFR and other selective media after 3 weeks in EGF-replete media, relative to control.

**Supplementary Figure 2. A)** Bar plot for the ratio of body weight to colon length in mice treated with DSS in water vs. water only control. Error bars represents standard deviation. n=3. **B)** Representative H&E sections of colons from a DSS-treated (right) and a control mouse (left).

**Supplementary Figure 3. A)** Targeting sites and sgRNA that were used to target *Apc* in normal mouse colonoids. **B)** Whole exome sequencing of *Apc*^mut^ colonoids confirms the presence of biallelic truncating mutations at the expected site of targeted CRISPR/Cas9 genome editing. **C)** Representative brightfield images of *Apc*^mut^ colonoids with characteristic spheroid morphology, as well as representative image from other colonoid lines. **D)** Representative brightfield images of colonoid lines corresponding to Day 0 and Day 7 of the selective challenges detailed in Figure 1B.

**Supplementary Figure 4.** The number of differentially expressed genes in iEGFR colonoids relative to control organoids, log_2_ fold changes ≥ 2 and Bonferroni *p*-value < 0.05. **A)** upregulated genes, **B)** downregulated genes.

**Supplementary Figure 5.** Chromosome copy number in control (euploid) and higher-passage iEGFR colonoids (~passage 54, aneuploid) quantified by qPCR, indicating loss of one copy of chromosome 13. n = 3 technical replicates.

**Supplementary Figure 6. A)** Representative confocal fluorescence images (Z sections with maximum projection) of colonoids stained with γH2AX antibodies (red) and DAPI (blue). Doxorubicin and No Ab (no antibody) represent the positive and negative controls, respectively. Scale bars = 50μm. **B)** The percentage of γH2AX positive cells/colonoid is represented as a box and whisker plot. Transverse lines represent the median, boxes show 25^th^-75^th^ percentile and the whiskers represent the lowest and highest values within 1.5 times the interquartile range. ns = not statistically significant, *p* ≥ 0.05 2-tailed, non-paired student t-test.

**Supplementary Table 1.** Unfiltered differentially expressed genes in iEGFR and DSS iEGFR colonoid lines vs. their respective controls.

**Supplementary Table 2.** Mouse primer sequences for chromosome karyotyping by qRT-PCR.

**Supplementary Table 3.** Mouse primer sequences for qRT-PCR validation of RNA-seq data

**Supplementary Video 1 (Control colonoids)**

Example of normal cell division.

**Supplementary Video 2 (iEGFR colonoids)**

Example of an erroneous division with a multipolar mitosis.

**Supplementary Video 3 (iEGFR colonoids)**

Example of an erroneous division with an anaphase bridge.

**Supplementary Video 4 (DSS control colonoids)**

Example of an erroneous division with a lagging chromosome.

**Supplementary Video 5 (DSS iEGFR colonoids)**

Example of an erroneous division with mitotic failure.

**Supplementary Video 6 (*Apc***^mut^ **colonoids)**

Example of lagging chromosomes.

## REFERENCES

1. Basak, O. et al. Induced Quiescence of Lgr5+ Stem Cells in Intestinal Organoids Enables Differentiation of Hormone-Producing Enteroendocrine Cells. Cell Stem Cell 20, 177–190.e4 (2017).

2. Muzny, D. M. et al. Comprehensive molecular characterization of human colon and rectal cancer. Nature 487, 330–337 (2012).

3. Fujii, M. et al. A Colorectal Tumor Organoid Library Demonstrates Progressive Loss of Niche Factor Requirements during Tumorigenesis. Cell Stem Cell 18, 827–838 (2016).

4. Clevers, H. The intestinal crypt, a prototype stem cell compartment. Cell 154, 274–84 (2013).

5. Powell, D. W., Pinchuk, I. V., Saada, J. I., Chen, X. & Mifflin, R. C. Mesenchymal Cells of the Intestinal Lamina Propria. Annu. Rev. Physiol. (2011) doi:10.1146/annurev.physiol.70.113006.100646.

6. Valenta, T. et al. Wnt Ligands Secreted by Subepithelial Mesenchymal Cells Are Essential for the Survival of Intestinal Stem Cells and Gut Homeostasis. Cell Rep. 15, 911–918 (2016).

7. Sato, T. et al. Long-term Expansion of Epithelial Organoids From Human Colon, Adenoma, Adenocarcinoma, and Barrett’s Epithelium. Gastroenterology 141, 1762–1772 (2011).

8. van de Wetering, M. et al. Prospective Derivation of a Living Organoid Biobank of Colorectal Cancer Patients. Cell 161, 933–945 (2015).

9. Kobayashi, H. et al. The balance of stromal BMP signaling mediated by GREM1 and ISLR drives colorectal carcinogenesis. Gastroenterology (2020) doi:10.1053/j.gastro.2020.11.011.

10. Drost, J. et al. Sequential cancer mutations in cultured human intestinal stem cells. Nature 521, 43–47 (2015).

11. Matano, M. et al. Modeling colorectal cancer using CRISPR-Cas9–mediated engineering of human intestinal organoids. Nat. Med. 21, 256–262 (2015).

12. Li, X. et al. Oncogenic transformation of diverse gastrointestinal tissues in primary organoid culture. Nat. Med. 20, 769–777 (2014).

13. Ritsma, L. et al. Intestinal crypt homeostasis revealed at single-stem-cell level by in vivo live imaging. Nature 507, 362–365 (2014).

14. Carroll, T. D. et al. Interkinetic nuclear migration and basal tethering facilitates post-mitotic daughter separation in intestinal organoids. J. Cell Sci. 130, 3862–3877 (2017).

15. Kinchen, J. et al. Structural Remodeling of the Human Colonic Mesenchyme in Inflammatory Bowel Disease. Cell 175, 372–386.e17 (2018).

16. Colotta, F., Allavena, P., Sica, A., Garlanda, C. & Mantovani, A. Cancer-related inflammation, the seventh hallmark of cancer: links to genetic instability. Carcinogenesis 30, 1073–1081 (2009).

17. Coussens, L. M. & Werb, Z. Inflammation and cancer. Nature 420, 860–867 (2002).

18. Holloway, E. M. et al. Mapping Development of the Human Intestinal Niche at Single-Cell Resolution. Cell Stem Cell (2020) doi:10.1016/j.stem.2020.11.008.

19. Fawkner-Corbett, D. et al. Spatiotemporal analysis of human intestinal development at single-cell resolution. Cell S009286742031686X (2021) doi:10.1016/j.cell.2020.12.016.

20. Ovid: Peptide growth factors in the intestine. http://ovidsp.dc2.ovid.com/ovid-b/ovidweb.cgi?T=JS&PAGE=fulltext&D=ovft&AN=00042737-200107000-00002&CHANNEL=CrossRef.

21. Eichele, D. D. & Kharbanda, K. K. Dextran sodium sulfate colitis murine model: An indispensable tool for advancing our understanding of inflammatory bowel diseases pathogenesis. World J. Gastroenterol. 23, 6016–6029 (2017).

22. Chassaing, B., Aitken, J. D., Malleshappa, M. & Vijay-Kumar, M. Dextran sulfate sodium (DSS)-induced colitis in mice. Curr. Protoc. Immunol. 104, 15.25.1–15.25.14 (2014).

23. Wirtz, S. et al. Chemically induced mouse models of acute and chronic intestinal inflammation. Nat. Protoc. 12, 1295–1309 (2017).

24. Sato, T. et al. Paneth cells constitute the niche for Lgr5 stem cells in intestinal crypts. Nature 469, 415–8 (2011).

25. Drost, J. et al. Sequential cancer mutations in cultured human intestinal stem cells. Nature 521, 43–47 (2015).

26. Schwank, G. & Clevers, H. CRISPR/Cas9-Mediated Genome Editing of Mouse Small Intestinal Organoids. in Gastrointestinal Physiology and Diseases: Methods and Protocols (ed. Ivanov, A. I.) 3–11 (Springer, 2016). doi:10.1007/978-1-4939-3603-8_1.

27. Dekkers, J. F. et al. High-resolution 3D imaging of fixed and cleared organoids. Nat. Protoc. 14, 1756–1771 (2019).

28. Bolhaqueiro, A. C. F. et al. Live imaging of cell division in 3D stem-cell organoid cultures. in (2018). doi:10.1016/bs.mcb.2018.03.016.

29. Miura, K. Temporal-Color Code (Version 101123) https://github.com/fiji/fiji/blob/master/plugins/Scripts/Image/Hyperstacks/Temporal-Color_Code.ijm (2010).

30. Pavelka, N. et al. Aneuploidy confers quantitative proteome changes and phenotypic variation in budding yeast. Nature 468, 321–325 (2010).

31. Dobin, A. et al. STAR: ultrafast universal RNA-seq aligner. Bioinforma. Oxf. Engl. 29, 15–21 (2013).

32. Liao, Y., Smyth, G. K. & Shi, W. The Subread aligner: fast, accurate and scalable read mapping by seed-and-vote. Nucleic Acids Res. 41, e108 (2013).

33. Love, M. I., Huber, W. & Anders, S. Moderated estimation of fold change and dispersion for RNA-seq data with DESeq2. Genome Biol. 15, 550 (2014).

34. Raudvere, U. et al. g:Profiler: a web server for functional enrichment analysis and conversions of gene lists (2019 update). Nucleic Acids Res. 47, W191–W198 (2019).

35. Kim, J. J., Shajib, M. S., Manocha, M. M. & Khan, W. I. Investigating intestinal inflammation in DSS-induced model of IBD. J. Vis. Exp. JoVE (2012) doi:10.3791/3678.

36. Nowak, M. A. et al. The role of chromosomal instability in tumor initiation. Proc. Natl. Acad. Sci. U. S. A. 99, 16226–16231 (2002).

37. Moniz, S. et al. Protein kinase WNK2 inhibits cell proliferation by negatively modulating the activation of MEK1/ERK1/2. Oncogene 26, 6071–6081 (2007).

38. James, R. G. et al. Bruton’s tyrosine kinase revealed as a negative regulator of Wnt-beta-catenin signaling. Sci. Signal. 2, ra25 (2009).

39. Yasuda, Y. et al. Human NINEIN polymorphism at codon 1111 is associated with the risk of colorectal cancer. Biomed. Rep. 13, 1–1 (2020).

40. Fischer, A. H. et al. The cytologic criteria of malignancy. J. Cell. Biochem. 110, 795–811 (2010).

41. Hanahan, D. & Weinberg, R. A. Hallmarks of cancer: The next generation. Cell vol. 144 (2011).

42. Rancati, G. et al. Aneuploidy Underlies Rapid Adaptive Evolution of Yeast Cells Deprived of a Conserved Cytokinesis Motor. Cell 135, 879–893 (2008).

43. Huch, M. & Koo, B.-K. Modeling mouse and human development using organoid cultures. Development 142, 3113–3125 (2015).

44. Rusan, N. M. & Peifer, M. Original CIN: reviewing roles for APC in chromosome instability. J. Cell Biol. 181, 719–726 (2008).

45. Killcoyne, S. et al. Genomic copy number predicts esophageal cancer years before transformation. Nat. Med. 26, 1726–1732 (2020).

46. Miroshnychenko, D. et al. Spontaneous cell fusions as a mechanism of parasexual recombination in tumour cell populations. Nat. Ecol. Evol. 1–13 (2021) doi:10.1038/s41559-020-01367-y.

47. Schwitalla, S. et al. Intestinal Tumorigenesis Initiated by Dedifferentiation and Acquisition of Stem-Cell-like Properties. Cell 152, 25–38 (2013).

48. Miyoshi, H. et al. Prostaglandin E2 promotes intestinal repair through an adaptive cellular response of the epithelium. EMBO J. 36, 5–24 (2017).

49. Yui, S. et al. YAP/TAZ-Dependent Reprogramming of Colonic Epithelium Links ECM Remodeling to Tissue Regeneration. Cell Stem Cell 22, 35–49.e7 (2018).

50. Dvorak, H. F. Tumors: Wounds That Do Not Heal—Redux. Cancer Immunol. Res. 3, 1–11 (2015).

51. Ge, Y. et al. Stem Cell Lineage Infidelity Drives Wound Repair and Cancer. Cell 169, 636–650.e14 (2017).

52. Losick, V. P. Wound-Induced Polyploidy Is Required for Tissue Repair. Adv. Wound Care 5, 271–278 (2016).

53. Lucchetta, E. M. & Ohlstein, B. Amitosis of Polyploid Cells Regenerates Functional Stem Cells in the Drosophila Intestine. Cell Stem Cell 20, 609–620.e6 (2017).

54. Tan, G., Huang, C., Chen, J. & Zhi, F. HMGB1 released from GSDME-mediated pyroptotic epithelial cells participates in the tumorigenesis of colitis-associated colorectal cancer through the ERK1/2 pathway. J. Hematol. Oncol.J Hematol Oncol 13, 149 (2020).

55. Xia, X. et al. The role of pyroptosis in cancer: pro-cancer or pro-“host”? Cell Death Dis. 10, 650 (2019).

56. Olafsson, S. et al. Somatic Evolution in Non-neoplastic IBD-Affected Colon. Cell S0092867420308138 (2020) doi:10.1016/j.cell.2020.06.036.

57. Nanki, K. et al. Somatic inflammatory gene mutations in human ulcerative colitis epithelium. Nature 577, 254–259 (2020).

58. Khan, K. et al. Targeting EGFR pathway in metastatic colorectal cancer-tumour heterogeniety and convergent evolution. Crit. Rev. Oncol. Hematol. 143, 153–163 (2019).

59. Lee-Six, H. et al. The landscape of somatic mutation in normal colorectal epithelial cells. Nature 574, 532–537 (2019).

60. Nicholson, A. M. et al. Fixation and Spread of Somatic Mutations in Adult Human Colonic Epithelium. Cell Stem Cell 22, 909–918.e8 (2018).

61. Smith, A. L. M. et al. Age-associated mitochondrial DNA mutations cause metabolic remodeling that contributes to accelerated intestinal tumorigenesis. Nat. Cancer 1, 976–989 (2020).

