## Supplementary Table 2 for "Colonic epithelial adaptation to EGFR-independent growth induces chromosomal instability and is accelerated by prior injury"

| **Name** | **Forward Primer** | **Reverse Primer** |
| --- | --- | --- |
| MMChr01 | ACAACACTCATCGCAGGCTAAGGA | TAATCAAAGCTGCCACAGCCACAC |
| MMChr02 | GGGATAGTGATGGAATGGGC | AGCCTTCATGTGCTCATCTG |
| MMChr03 | TCAGAACTTAAAATACCAAAACCAGG | CTCAGTTCAAGAGTTAGTCTCAGAG |
| MMChr04 | GATGCCAGGAGACGAGAATAC | CACAATAGACAGTGCCTACCAG |
| MMChr05 | AGGTGTCTTACAGGAATATGAACTG | TCCATTTACATGACCTAGCCAC |
| MMChr06 | GGAAGCATGTGAAAGGTGTTG | TCCTTGCATGCCAATATACCC |
| MMChr07 | TGCCCTGTTCTGATGGGATTCTCT | AAAGGGAGCTAGCTGGCTGAAGAA |
| MMChr08 | TGGGCTTGAATTTGGGAGG | GAGAGGGAGTGGCTGATTTG |
| MMChr09 | ACGGAAACTCAACCTGGAATG | GCTTCGGTGATACAGATCCAG |
| MMChr10 | CAGCTCTGGGTTCAGATGAAG | TCCTGTCAGTACTTCAGCAATG |
| MMChr11 | ACCACCGCCCAGTCATCTTTCATA | AGCACTTGACACATGTCCTCCCTT |
| MMChr12 | GATGGTCTTGGGCTACACG | GCCTTCATCCTGCTTGAGTG |
| MMChr13 | CTTTGCGGATGAGGAAAACG | TGTGTCTCCTGCTTTGCTAC |
| MMChr14 | TCTACACGGCAGGGATTTG | ACAGAGTTTACAGAAGCTGGG |
| MMChr15 | CAGTATGCTCGTTCTGGAGTG | TCCTTCCCTTACTCTACCATCC |
| MMChr16 | TGGACACAGGAAGATAGCATTG | AAGGGCACATCTTCCATGAG |
| MMChr17 | GCCTGGAAGTTGCTATGATTG | TGCTGAACTTGAGGTGTTACC |
| MMChr18 | TTCAGGCCGTGATTAACTGG | CATCTTGTGCTCCCTCTACTG |
| MMChr19 | TTGACTTCGCCACTGTATCC | GACGAACAGAATGTCCACAATG |
| MMChrX | ATGCCATGAAATGTTGAATCCAG | TCTTCCTCTCTACCACCTTCTTC |
| MMChrY | AACAATACCCATCGCCTGG | AGGACACTGATTCACATGGAC |

**Supplementary Table 2.** Primer sequences for chromosome karyotyping by qRT-PCR.
