## Supplementary Table 3 for "Colonic epithelial adaptation to EGFR-independent growth induces chromosomal instability and is accelerated by prior injury"

**Supplementary Table 3.** Primer sequences used for qRT-PCR.

| **Gene** | **Forward** | **Reverse** |
| --- | --- | --- |
| ***Igfbp7*** | AGAAGGCCATCACCCAGGTC | GGGCGTCACTATGGAAGGAC |
| ***Mycn*** | CCGGAGAGGATACCTTGAGC | GAGGTCTTGGGACGCACAGT |
| ***Ninl*** | TAGAGCCCTCCGATGAGGAA | GTTGCAGTGGCAGAGTCACC |
| ***Wwtr1*** | CACAGCAGCATGCACATCTC | TCATCACCTTCCTGGGGTCT |
| ***Gsdmcl2*** | ACAGCCTCAGCAGCTCACAG | TTCACCTCTGGGTCAGGACA |
| ***Muc1*** | TCTCTGGAAGACCCCAGCTC | AAGTACCCTCCCGGAAAACC |
| ***Clu*** | AGATTCAGAACGCCGTCCA | GGGAATGCCTTCAGCTTCAT |
| ***Itga1*** | GCGGGACTCCTGCTGTTAAT | GATTGAGGCAAACCTGAGGA |
| ***Gpc1*** | TTGCCGAAATGTGCTCAAAG | CACACCGCCAATGACACTCT |
| ***Ppp1r3d*** | TGCTGAAGGTACCGAGGACA | GTCGCGACCATCATTGTTGT |
| ***Cav1*** | ATGGCAGACGAGGTGACTGA | TGTCCCTTCTGGTTCTGCAA |
| ***Dsg3*** | GCCCTGGGACAGGATGTAGA | TCTGCATCCGTGGCATTTAG |
| ***Il5ra*** | CCCAAATCCTGACCAAGAGC | ACGCTAGCTGCAAAGCCTTC |
| ***Efemp1*** | AGGTCCGTGCCTTCAGACAT | TGCACTCACAGGGCTTGTTT |
| ***Hmcn1*** | CTTTGGTGCCCTTCCATGAT | TAGCCCCGATGCTCATTTCT |
| ***Hprt*** | GTCCCAGCGTCGTGATTAGC | GGCCTCCCATCTCCTTCATGA |
| ***ActB*** | ATGCTCCCCGGGCTGTATTC | CGTCTCCGGAGTCCATCACA |
| ***Gsdmc2*** | CCTTTGAGAGTTCAGCCAGTT | GGCTTGTGGATAAATGACATCG |
| ***Gsdmc3*** | TGAATACTTTTCCGACCTTCCC | CAAGGATACTATGCTGACAGGA |
| **Gsdmc4** | CCTCAGATCTCTTCCTTGAAGC | CCTTTGTGAATAGTCTGCCATGA |
