## Supplementary Figures 1-6 for "Colonic epithelial adaptation to EGFR-independent growth induces chromosomal instability and is accelerated by prior injury"

**Supplementary Figure 1**

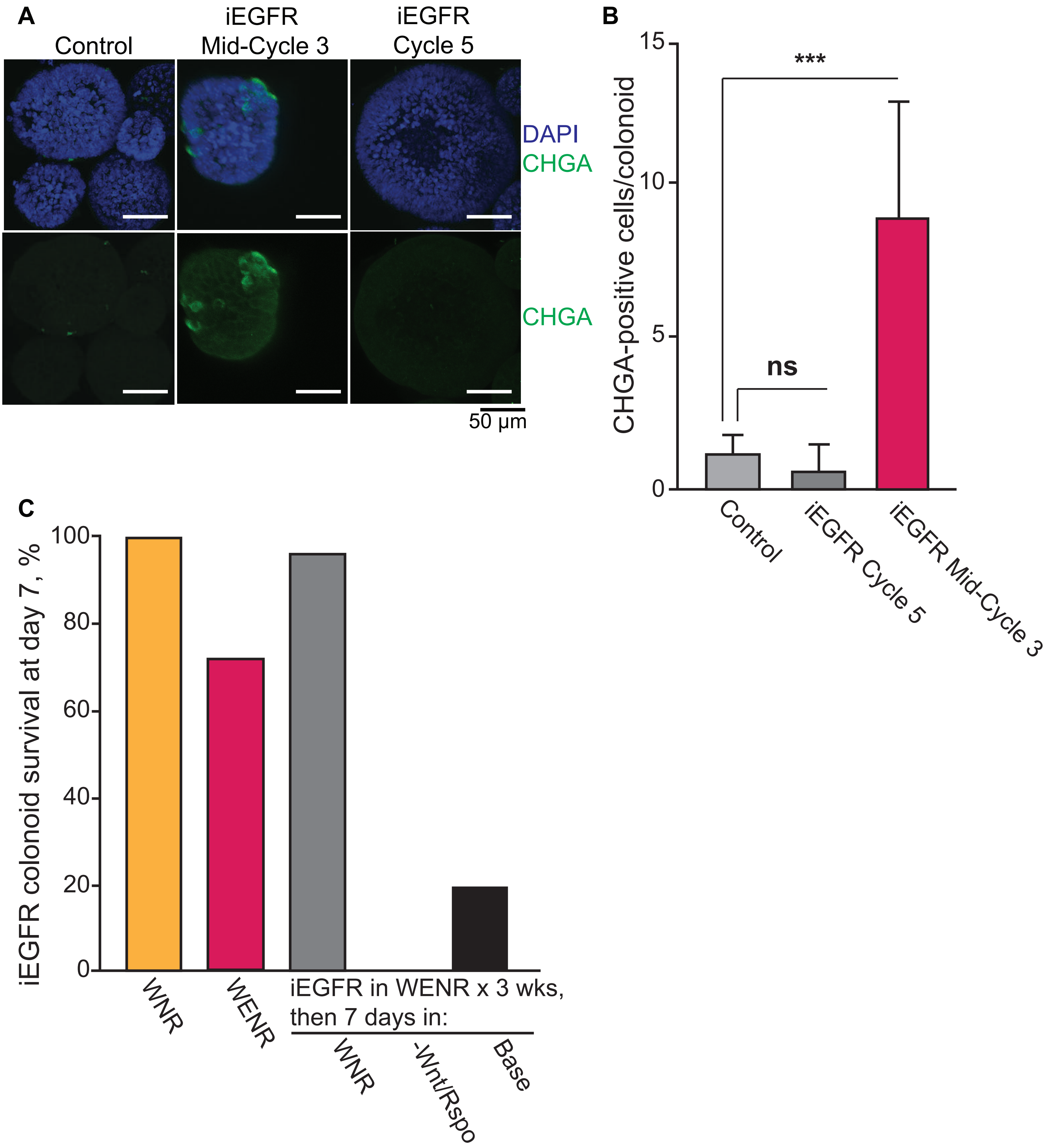

**Supplementary Figure 2**

**A**

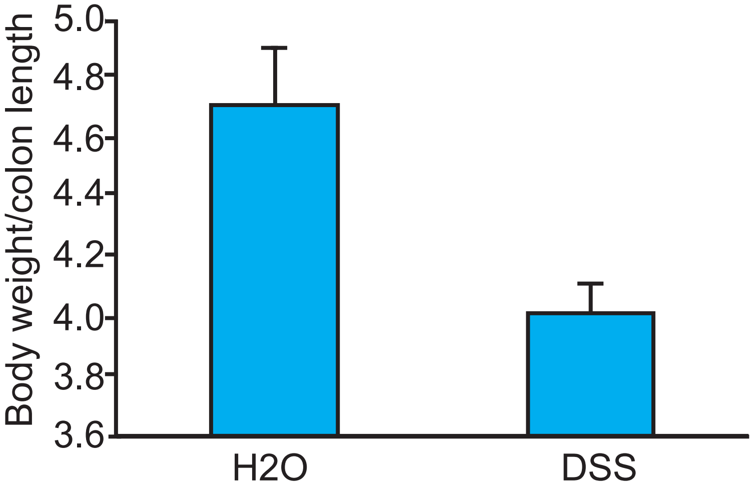


**B**

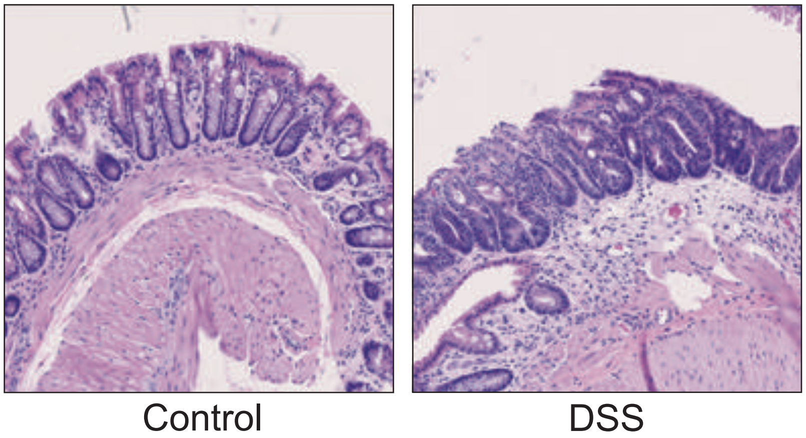

### Supplemental Figure 3

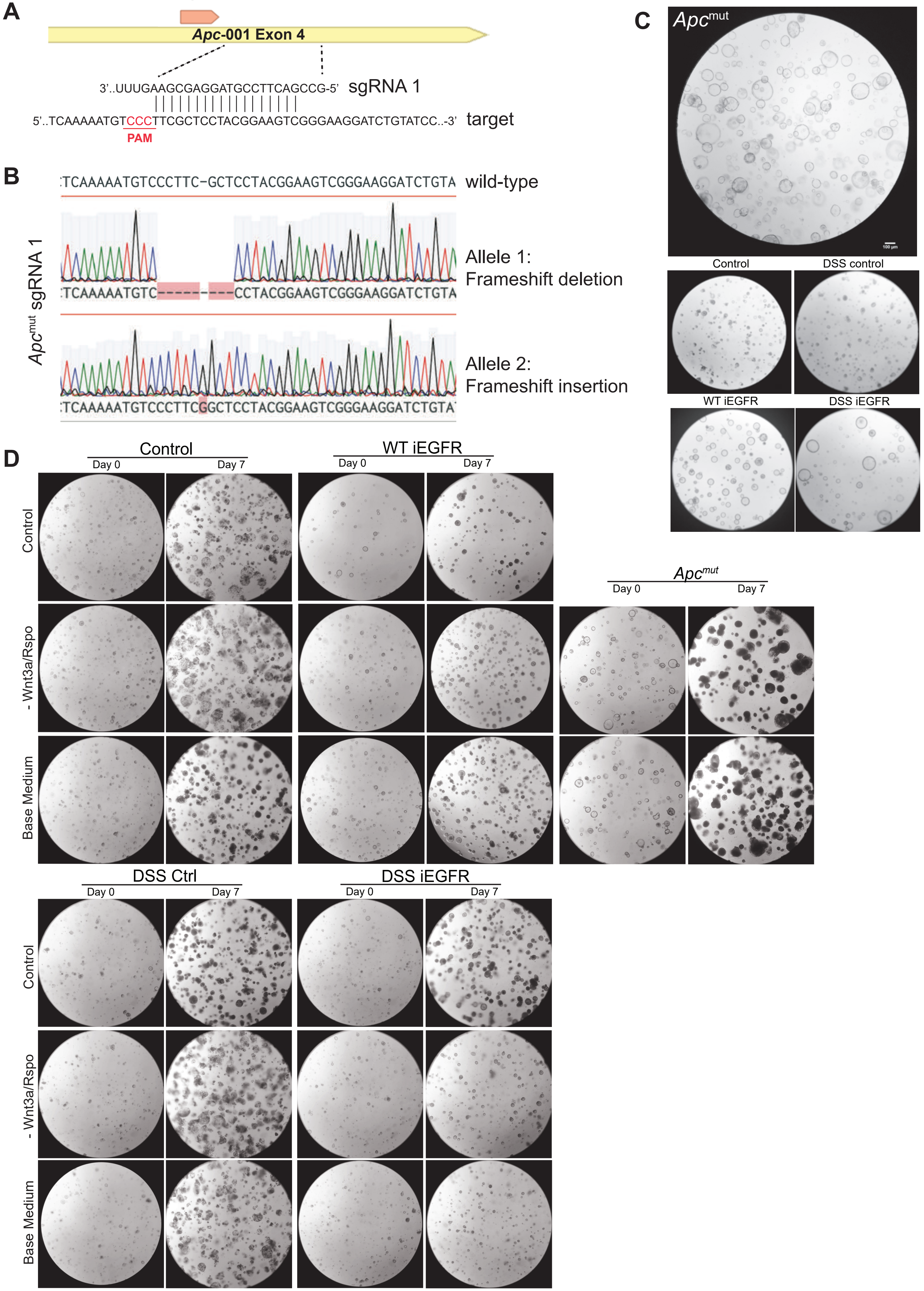

**Supplementary Figure 4**

A

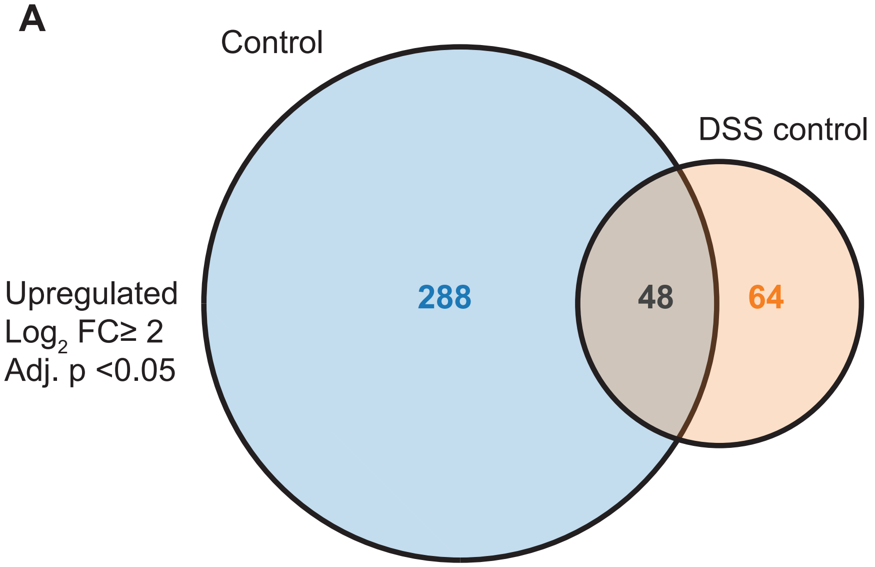


B

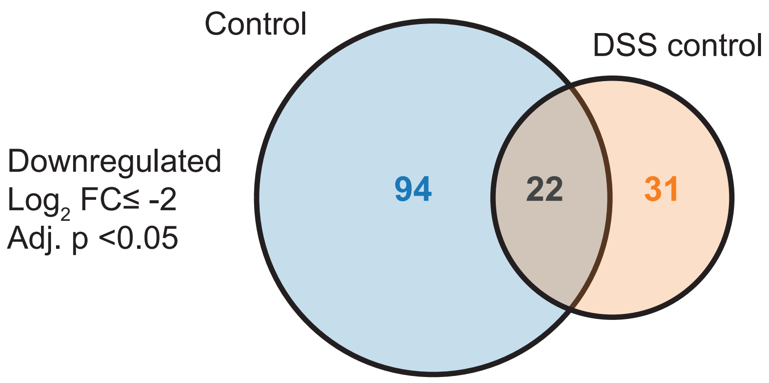

### Supplementary Figure 5

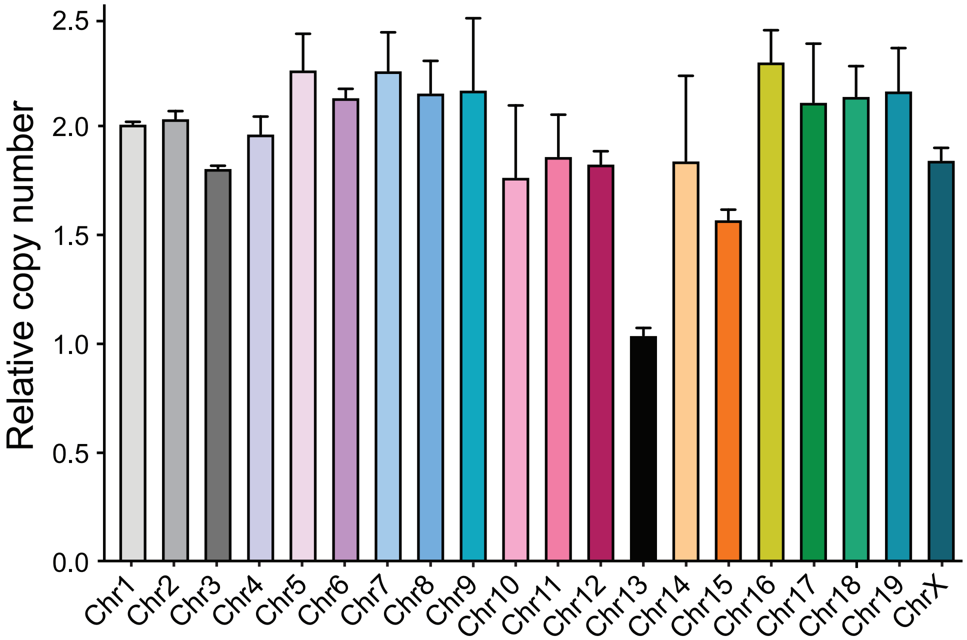

**Supplementary Figure 6**

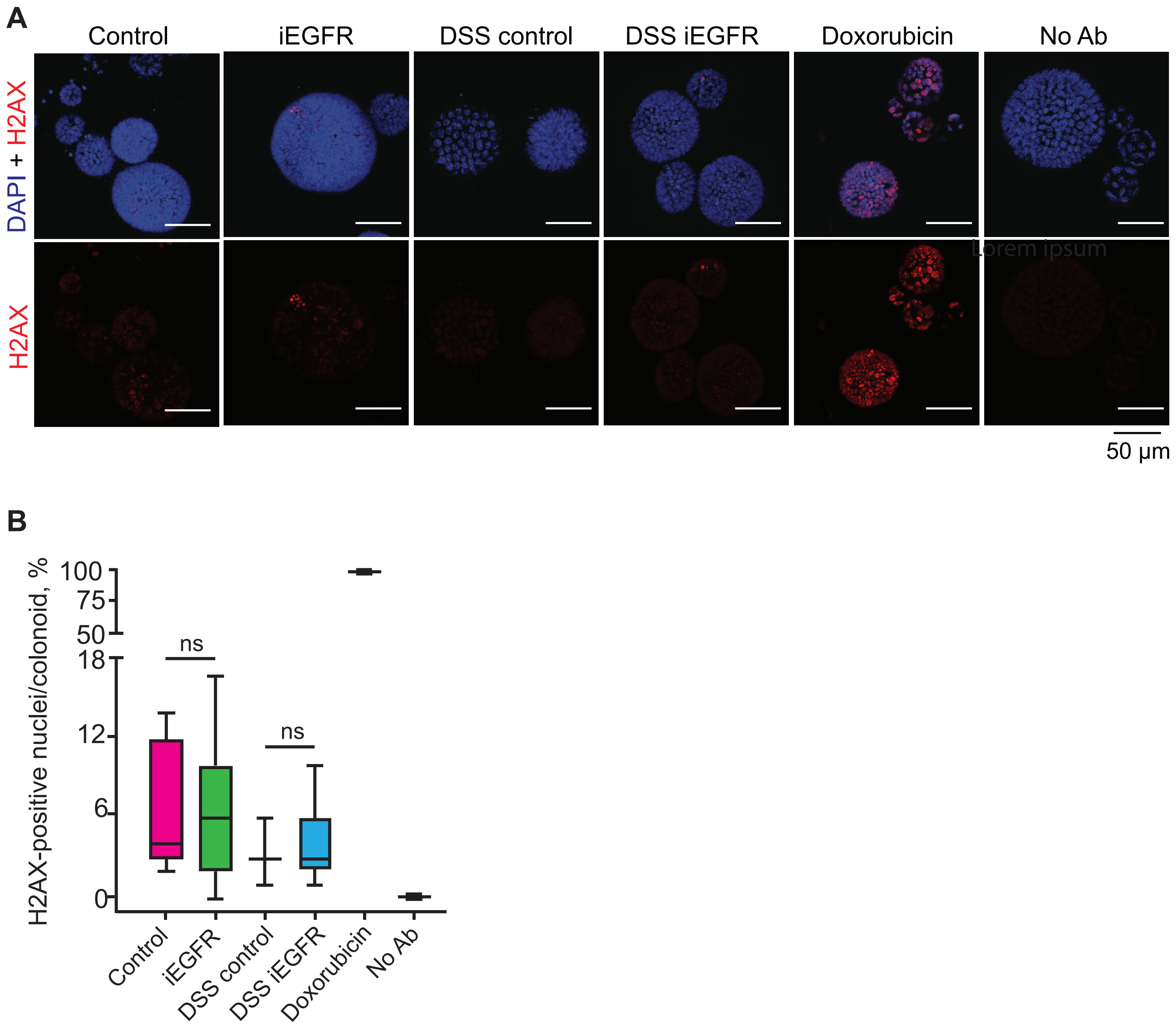
